# Plasticity and Artificial Selection for Developmental Mode in a Poecilogonous Sea Slug

**DOI:** 10.1101/2020.03.06.981324

**Authors:** Serena A. Caplins

## Abstract

Developmental mode describes the means by which larvae are provisioned with the nutrients they need to proceed through development and typically results in a trade-off between offspring size and number. The sacoglossan sea slug *Alderia willowi* exhibits intraspecific variation for developmental mode (= poecilogony) that is environmentally modulated with populations producing more yolk-feeding (lecithotrophic) larvae during the summer, and more planktonic feeding (planktotrophic) larvae in the winter. I found significant family level variation in the reaction norms between 17 maternal families of *A. willowi* when reared in low (16 ppt) versus high (32 ppt) salinity. I documented a significant response to selection for lecithotrophic larvae, the proportion of which increased 32% after three generations of selection in high salinity, and 18% after 2 generations in low salinity (realized heritability: 0.365 ± 0.024). The slope of the reaction norm was maintained following one generation of selection for lecithotrophy and one generation of selfing. The rapid response to selection favoring one developmental mode may speak to the rarity of intraspecific variation for developmental mode, which could fix for one mode over another much more readily than has generally been assumed from studies of less plastic organisms.

## Introduction

Marine invertebrates exhibit astonishing levels of morphological diversity in their adult forms. Their larvae, however, can be broadly grouped into a few developmental modes that while also morphologically variable, share many functional similarities within and between phyla (Thorson 1950; Strathmann 1978, Collin and Moran 2017). The inferred ancestral state for many phyla is planktotrophic development, involving the production of many relatively small larvae that feed in the plankton prior to settlement and metamorphosis to the adult form (Strathmann 1978; McHugh and Rouse 1998). Lecithotrophy (non-feeding) is the most common alternative to planktotrophic development, in which a few relatively large larvae contain substantial amounts of yolk, such that they do not need to feed in the plankton before metamorphosis (McEdward and Janies 1997; Marshall et al 2017). Both planktotrophy and lecithotrophy are found in all invertebrate bilaterian phyla and are widespread phylogenetically (Ruppert 2004). Numerous studies suggest that the evolution of developmental mode is primarily uni-directional (from planktotrophy to lecithotrophy), though reversions or convergent re-evolution of planktotrophy from lecithotrophy are possible (Marshall, Raff, and Raff 1994; Kupriyanova, Macdonald, and Rouse 2006; Collin and Miglietta 2008). Modifications to developmental mode can evolve rapidly (< 20 kya, Puritz et al. 2012), and frequently in marine invertebrates with closely related sister species often displaying alternate modes of development (Krug et al. 2015; Jeffery, Emlet, and Littlewood 2003). These patterns have intrigued evolutionary biologists for decades, as we ask how does lecithotrophy evolve? Are there many or few genetic changes in the evolution of lecithotrophy? Are there environmental factors that select for one developmental mode over another?

Phenotypic plasticity, the ability of a single genotype to produce multiple phenotypes depending on the environment, is a common adaptation to spatial or temporal environmental heterogeneity (Via et al. 1995). Yet, the evolutionary role of plasticity is highly context-dependent, sometimes fueling evolution by moving the mean phenotype in the direction favored by selection, and other times hindering evolution through the lack of a genetic response to selection on a variable phenotype (see Pfennig et al. 2010). Within a generation, phenotypic variation can shift after a selective event (e.g., a sudden change in environment, or predation), but the *response* to selection in the following generation reveals whether there is sufficient genetic variation underlying phenotypic variation for evolution to occur (Falconer 1960). Selection experiments can thus reveal the extent to which there is heritable genetic variation for a plastic phenotype and are a powerful means of exploring the potential for adaptive evolution under highly controlled environmental conditions, simplifying the study of environmentally influenced quantitative traits (Scheiner 2002; Fuller, Baer, and Travis 2005). Selection experiments can also reveal the effect of specific environmental factors, which may influence the response to selection by either revealing or masking ‘cryptic’ genetic variants (Falconer 1960; Paaby and Rockman 2014).

In this paper, I examine the role of environmentally modulated plasticity in the evolution of larval developmental mode in a marine gastropod that exhibits phenotypic plasticity for developmental mode. Species that are polymorphic for the type of larvae they produce provide a novel means of addressing the evolution of macro-evolutionary patterns in a micro-evolutionary framework. This polymorphism, termed *poecilogony*, occurs when a single species produces both planktotrophic and lecithotrophic larvae (Knott and McHugh 2012). Poecilogony typically manifests as either variation for developmental mode within a population, or variation between populations (Collin 2012). Poecilogonous species provide a clear way to explore the selection pressures and corresponding genetic and transcriptomic changes that drive the evolution of developmental mode. Poecilogonous species are key to understanding the *minimum number* of genetic changes that underlie developmental mode, as they provide a view into the specific alternative developmental pathways that give rise to these modes in the absence of interspecific differences (Alan and Pernet 2007, Knott and McHugh 2012; Zakas et al. 2018).

There is only one poecilogonous species that exhibits environmental modulation in its expression of developmental mode, the sea slug *Alderia willowi*, where egg size and number are negatively correlated and bimodally distributed, with individual clutches consisting of either many small eggs that develop into planktotrophic larvae, or relatively few large eggs that can successfully metamorphose into juvenile slugs without feeding (Krug 1998, 2001). The relative frequency of clutches containing either planktotrophically or lecithotrophically developing eggs varies seasonally (Ellingson and Krug 2016) and in response to starvation (Krug 1998) or lab conditions (Krug 2007). In field populations, individuals produce a greater proportion of clutches containing a few large eggs with lecithotrophic development during the summer months, and more clutches with many small, planktotrophically developing eggs in the winter (Krug et al. 2012). Lab populations show a similar pattern, with conditions that mimic summer resulting in more lecithotrophically developing embryos (Krug et al. 2012). Despite these seasonal trends that are consistent across years, lab and field populations exhibit variation in the proportion of lecithotrophic clutches produced seasonally and geographically (Patrick J. Krug et al. 2007; Patrick J. Krug, Gordon, and Romero 2012). Herein I examine the extent of genetic variation and environmental influence on developmental mode in maternal families of *A. willowi*. I measured the response to selection for lecithotrophy and evaluate whether one generation of selection affects the direction or degree of plasticity.

## Methods

### Study system

An *Alderia willowi* egg mass consists of dozens to hundreds of eggs strung together and surrounded by a thick jelly like substance (Figure 1A). Each individual egg is surrounded by a transparent capsule the diameter of which scales closely with egg diameter (Figure 1B-D). In *A. willowi*, egg size is correlated with developmental mode large eggs (Mean ± SD: 105 ± 5 µm) develop into lecithotrophic larvae that metamorphose into juvenile slugs in ∼5 days, whereas small eggs (Mean ± SD: 68 ± 4 µm) give rise to planktotrophic larvae that only become metamorphically competent after 30 days of feeding on planktonic algae (Krug 1998). Both size classes of larvae can feed on phytoplankton, but the larger, lecithotrophically developing larvae do not need to feed to complete metamorphosis and occasionally develop into the juvenile stage while still encapsulated in their egg capsule, by-passing a swimming stage altogether (Krug 2001; Botello and Krug 2006). Infrequently, individual *A. willowi* produce mixed-egg clutches containing both lecithotrophic and planktotrophic embryos (Krug 1998). In these egg-masses, larvae with a larval shell diameter > 160 µm exhibit lecithotrophic development, whereas smaller larvae are all planktotrophic (Krug 1998).

**Figure 1.**
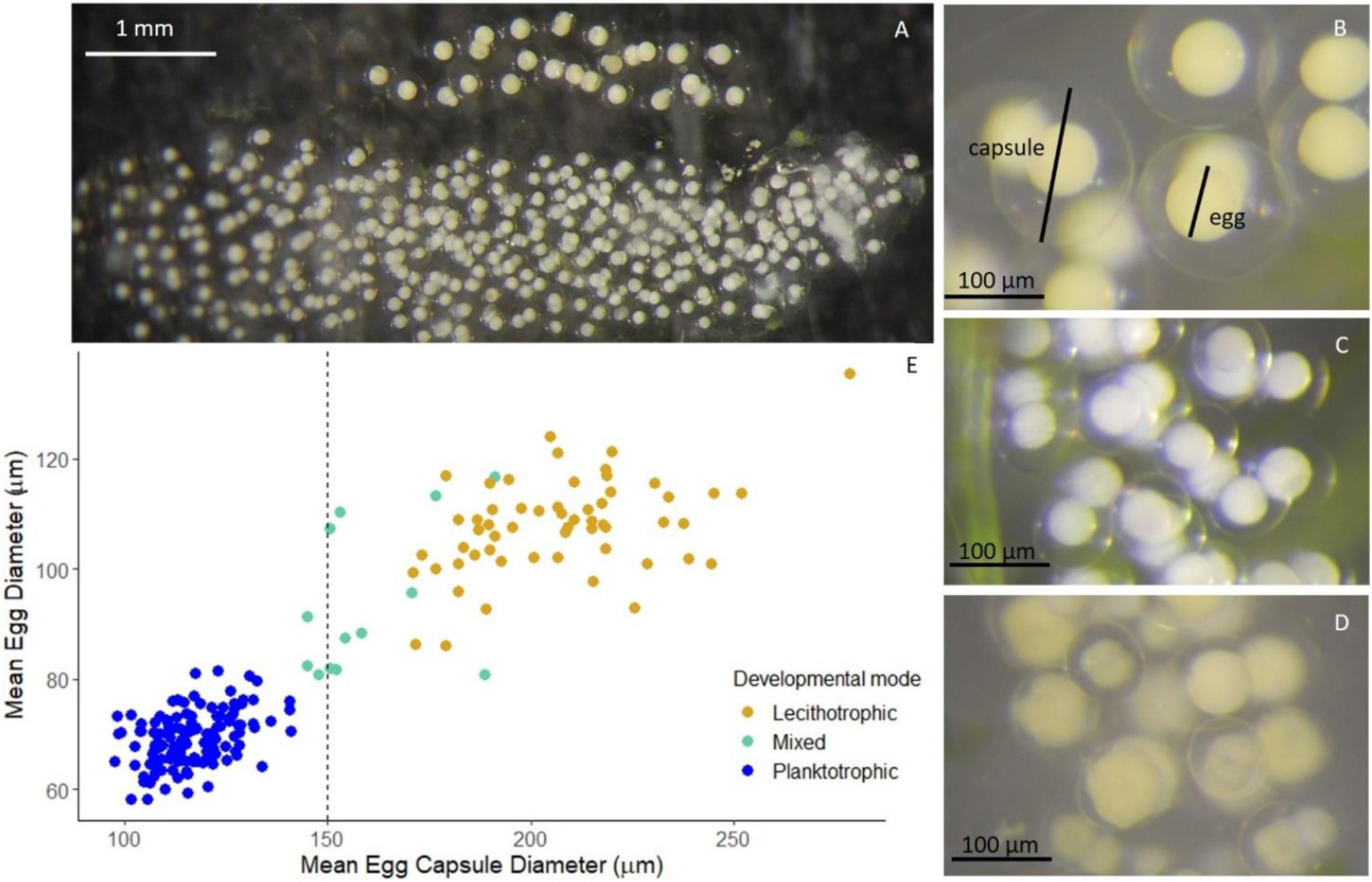
The upper egg mass in A) is a lecithotrophic egg mass, below which is a planktotrophic egg mass (10X magnification). Egg masses at increased magnification (50X) B) Lecithotrophic, C) Planktotrophic, and D) Mixed (planktotrophic and lecithotrophic, pictured in the gastrula stage) egg masses. A scatterplot of the relationship between mean egg diameter and mean egg capsule diameter calculated per egg mass is shown in E for a subset of the total egg masses measured in this study (i.e., 195 egg masses from 123 individuals).

Populations of *A. willowi* are found on mudflats in estuarine environments and can be extremely variable in density, from several dozen individuals/m^2^ to 1300 slugs/m^2^ (Garchow 2010). Individuals are typically polyandrous, with multiple matings via hypodermic insemination (Smolensky, Romero, and Krug 2009). At low densities, however, *A. willowi* exhibits “delayed selfing” (Smolensky, Romero, and Krug 2009). Self-fertilized egg masses are occasionally incompletely fertilized and *A. willowi* will continually deposit unfertilized or partially fertilized egg masses when reared in isolation (personal obvs., Smolensky, Romero, and Krug 2009).

### Effect of salinity on developmental mode

Adult *Alderia willowi* were collected from three sites in CA: Tomales Bay (20 June 2017 [38°06’59”N 122°51’16”W], Mill Valley (12 September 2017 [37°52’55”N 122°31’03”W] and Long Beach (14 September 2017, provided by Patrick Krug [33.73 N, 118.203 W]). I selected lecithotrophic egg masses from 17 adult slugs (Tomales Bay: 7; Mill Valley: 5; Long Beach: 5), from which the embryos constituted 17 maternal families consisting of unknown mixtures of full- or half-sibs. Embryos from each family were hatched and reared to the newly metamorphosed juvenile stage in 32 ppt filtered sea water at room temperature. Embryos were not provided with planktonic algae, and instead relied on their yolk stores to complete development. Once slugs were at the crawling juvenile stage, I haphazardly selected individuals from each maternal family to either the low (16 ppt) or high (32 ppt) salinity treatment. I placed 24-36 juvenile slugs in each salinity treatment per maternal family. Slugs were reared individually in 12-well culture dishes with a 5 ml volume per well and on a 14L:10D light cycle at room temperature (22 ± 2 °C). I covered each culture plate with plastic wrap that has a water-resistant adhesive on one side to keep slugs in their respective wells. Three times weekly, I fed slugs freshly field collected algae (*Vaucheria longicaulis*), carried out 50% water changes, and checked for newly deposited egg masses. Once slugs reached maturity and began laying egg masses, I photographed each egg mass using a Nikon CoolPix P7100 on a Wild Heerbrugg dissecting microscope at 50X magnification. I measured the diameter of three to six egg capsules which surround each individual egg in Image J (v1.52). For every egg mass that contained eggs that had yet to cleave, I measured the diameter of three to six eggs in addition to measuring the egg-capsule diameter.

As egg size can only be accurately measured prior to embryonic cleavage, and thus within the first 1-2 hours post oviposition, I used egg-capsule size as a proxy for developmental mode. I categorized developmental mode according to egg capsule size in a clutch/egg mass, assuming that egg capsules ≥ 150 µm develop lecithotrophically (Krug 1998). I used an R script to classify which egg masses were “mixed” based on egg capsule measurements. I verified these “mixed” egg masses through examination of the egg mass images. To explore the relationship between egg diameter and egg capsule diameter I plotted the mean per egg mass of egg diameter against the mean per egg mass of egg capsule diameter (Figure 1).

### Analysis of genetic variance and heritability

Models of quantitative genetics use population pedigree information to estimate genetic variance and heritability. Standard models of quantitative genetics assume traits have normal distributions; however, many traits are non-normally distributed (Hadfield and Others 2010). Generalized linear mixed models (GLMM) make use of a latent variable (ℓ) rather than the observed response, and in simulated data provide a better fit for binary traits than parent-offspring regression (de Villemereuil 2012). The latent variable of GLMMs incorporates non-normal trait distributions in quantitative genetics models. In this paper, the model takes the form below for each individual (i):

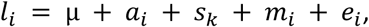

where µ is the mean phenotype in the population, *a* is the breeding value, *s* is the effect of salinity (on k levels, low or high), *m* is the maternal effect of each maternal family, and *e* is the error term. Salinity and maternal effects are input in the model as random effects.

I tested the effect of salinity on egg-mass type with the response either as a continuous variable with a gaussian distribution for egg-capsule size, or as a binary variable for developmental mode (e.g., lecithotrophy = 1, planktotrophy = 0). For the binary analysis, I assumed that egg-capsules > 150 µm contained eggs that developed lecithotrophically, and those capsules < 150 µm developed planktotrophically and was likewise analyzed as a threshold trait (i.e., a quantitative trait with discrete expression, see Roff 1994). Because the binary trait (developmental mode) is derived from the continuous trait (egg capsule size), I performed each analysis separately and not as a multivariate analysis. A GLMM requires a probit link-function to go from the latent Gaussian variable to the observed response variable. In the case of a threshold response this takes the form: P(y_i_ ^0,1^ = 1) = probit^-1^(ℓ_i_). The link function for the response of egg capsule size took the standard form for a gaussian response variable (see de Villemereuil 2018). The models were run in the R package *MCMCglmm* (Hadfield 2010). I specified priors for the gaussian model (egg capsule size) as a normal distribution with mean = zero and a small variance (1), and for the threshold model as a normal distribution with mean=zero and a large variance (1000), as described in de Villemereuil (2018). All analyses were performed in the R environment (v3.5.1) and the code used along with all the data presented in this paper are available on github (SerenaCaplins/GXE_A.willowi).

### Broad sense heritability

Heritability for threshold traits can be measured on two scales, the observed non-normally distributed phenotypic scale, and the normally distributed un-observed *liability* (Falconer 1960, de Villemereuil 2018). I used the R package *QGglmm* (de Villemereuil 2018) to calculate heritability on both the observed and liability scale for developmental mode, and on just the observed scale for egg-capsule diameter. I analyzed developmental mode as a binary trait (lecithotrophy = 1, planktotrophic = 0) in an MCMCglmm model set for a “threshold” distribution for 500,000 iterations with a burn-in phase of 3000 and a thinning interval of 10.

### Genetic correlation

Falconer (1960) noted that a phenotype produced in two environments could be viewed as two separate phenotypes, and thus a genetic correlation can be calculated between the two. This correlation can be used to determine the degree to which a phenotypic response is influenced by the environment, where a perfect correlation (= 1) between environments indicates zero environmental influence. This correlation also provides a prediction for how a given phenotype may respond to selection in a given environment (Falconer 1952). I used the family level proportion of lecithotrophic egg-masses produced in low and high salinity to evaluate the genetic correlation between salinities (see Via 1984; Roff 1996).

### Selection for lecithotrophy in low and high salinity

To evaluate the response to selection for lecithotrophy, I selected egg masses containing large eggs for three generations. The larvae from selected egg masses were at no time fed planktonic algae, and thus all that survived to the juvenile stage were lecithotrophic in their development. I reared slugs in their respective maternal salinities (low or high) each generation. The S_1_ and S_2_ generations were the product of self-fertilization, because the hermaphroditic slugs were raised in isolation. Slugs were reared in 12-well cell-culture plates, which were covered with plastic wrap as described above. I fed adult and juvenile slugs *V. longicaulis* and changed their water three times weekly. The parental generation uses data pooled from two experiments with 189 slugs from 17 families in low and 244 slugs from 17 families in high salinity. Data for the S1 and S2 generations were all collected concurrently using slugs from the parental generation that were collected from Long Beach and Mill Valley. A selection experiment was no performed on slugs from Tomales Bay and thus only data for a parental generation exists for this site. I measured egg capsule size for five capsules per egg mass in ImageJ (v1.52a). I calculated realized heritability on developmental mode using the breeder’s equation (R = h^2^S) modified for a threshold response using a probit transformation to translate the proportion of individuals expressing the trait of interest to a mean value for that trait (Walsh and Lynch 2018).

### Selection and the reaction norm

To evaluate whether the slope of the reaction norm changes following selection lecithotrophy, I reared slugs from the S_1_ generation from two of the sites mentioned above (Mill Valley and Long Beach) in either their maternal salinity or the alternate high or low salinity. Mill Valley and Long Beach are the northern and central range sites for *A. willowi*, respectively, and may be different in their response to salinity due to differences in seasonal annual rainfall (Garchow 2010). Fifty percent of every clutch was reared in either high or low salinity, as described previously. I measured egg capsule size for three to six egg capsules per egg mass in ImageJ (v1.52a). I tested the significance of the parental reaction norm against the reaction norm after selection using a linear model with the response egg capsule diameter against the predictors salinity and generation (pre or post selection).

## Results

### Effect of salinity on developmental mode

Egg capsule size closely predicts egg size, and egg size is a proxy for developmental mode (Figure 1, Krug 1998). Egg capsule size and egg size can be measured on a continuous scale, but both are bimodally distributed as shown in measurements of the mean egg diameter and mean egg capsule diameter per egg mass for 195 egg masses from 123 individuals in low (N = 70) and high (N = 125) salinity (Figure 1E). Egg size has a smaller standard deviation than egg capsule size (egg diameter SD = 0.017, egg capsule diameter SD = 0.043). Egg capsule size remains constant throughout development (Figure S1). Slugs began to deposit egg masses when they were an average of 17.5 days old. A total of 1958 egg masses were laid by 433 slugs from 17 families. The number of egg masses an individual laid ranged widely (mean =4.5, min = 0, max = 33). Most egg capsules within an egg mass were similar in size (mean = 136 µm, SD = 0.0097, and the two highest modes = 113 µm, and 182 µm). Fewer egg masses were laid in low salinity than in high salinity (1037 versus 1809, respectively). Proportionally there were fewer lecithotrophic egg masses produced in low salinity (17.6%) than in high (25.7%). Mixed egg masses contain both large eggs (> 150 µm capsule size) and small eggs (< 150 µm capsule size). Mixed egg masses occurred in both high and low salinities at similar proportions (6.3% in low salinity, 6.7% in high salinity). There was an effect of collection site or collection month, with the slugs collected in Tomales bay in June producing significantly fewer lecithotrophically developing egg masses (5%) than either the Long Beach (24%) or Mill Valley (22%) populations, both of which were collected in September (Figure 2A; linear model (egg capsule diameter ∼ region + salinity + eggmass type), region p-value < 0.001, salinity p-value = 0.0215, eggmass type p-value < 0.001, r^2^=0.76).

**Figure 2.**
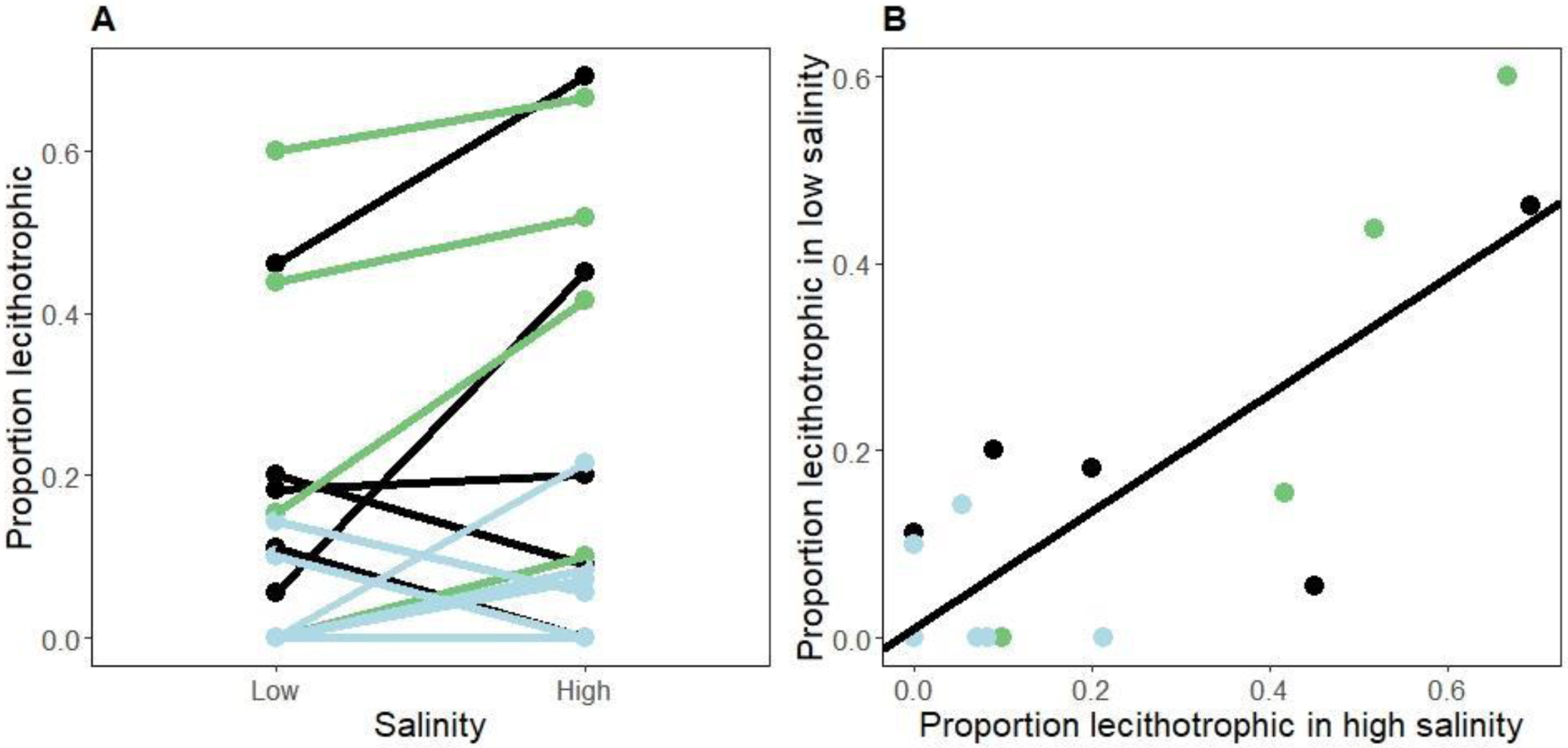
A) Family reaction norm for the proportion of lecithotrophy in low (16 ppt) and high (32 ppt) salinity and (B) the correlation between families for the proportion of lecithotrophy produced in low and high salinity (slope = 0.63, intercept = 0.001). Each line in A and each dot in B are a maternal family. Colors denote sampling sites: Tomales Bay, CA in green, Mill Valley, CA in light blue, and Long Beach, CA in black.

The reaction norm of the proportion of lecithotrophic egg masses reveals considerable variation for egg-mass type within and between families (Figure 2A). Most families show an increase in egg capsule size and in the proportion of lecithotrophic eggs in high salinity, although four families produced more lecithotrophic egg masses in low salinity than in high (Figure 2A, showing proportion lecithotrophic). Five families produced lecithotrophic egg masses in high salinity, but not in low salinity (Figure 2A, B). Likewise, one family produced no lecithotrophic eggs in either salinity, but also had the lowest survival rate in lab conditions of any other family. Three of the four reaction norms with negative slopes had a small sample size (N < 10). Offspring survival to adulthood was lower in low salinity than in high salinity (63% versus 81%, respectively). While survival declined in low salinity, survival was not significantly correlated with the proportion of lecithotrophy in either low or high salinity (linear model, low salinity r^2^ = 0.001, p = 0.89; high salinity r^2^ = 0.005, p = 0.77).

### Genetic correlations between environments

The family response to salinity is positively correlated across salinity treatments (Figure 2B; slope = 0.63; Y-intercept = 0.001, multiple r^2^ = 0.89, p-value 7.13e-05). This slope predicts the expected response to selection for developmental mode between high and low salinity: for every one-unit change in response to selection in high salinity, a corresponding 63% change should occur in low salinity.

### Analysis of genetic variance and heritability

The residuals for egg-capsule size against maternal family and salinity are bimodally distributed, as are the raw data (Figure S1). The analysis revealed a significant effect of salinity and family on egg-capsule size (MCMCglmm for Gaussian trait; salinity p-value = 0.004, maternal family p-value < 0.001). Broad sense heritability for egg capsule size was 0.54 (Table 1). The model testing the effect of salinity and maternal family on the proportion of lecithotrophic egg masses (the threshold model) also revealed a significant effect of salinity and maternal family on developmental mode (salinity p-value = 0.000644, maternal family p-value < 2e-05). Broad sense heritability on the observed scale was 0.23 and on the latent scale was 0.46 (Table 1). For both models, I assessed model fit by confirming that the effective sample size exceeded 1000, and the trace and density plots showed adequate mixing.

**Table 1.** Summary of model values and broad sense heritability where V_a_ is the additive genetic variance, and V_r_ is the link variance as computed in each model and corresponds to phenotypic variance.

| trait | distribution | $\mu$ | $V_a$ | $V_r$ | H<sup>2</sup> latent | H<sup>2</sup> obs. |
| --- | --- | --- | --- | --- | --- | --- |
| egg-capsule size | gaussian | 0.13 | 0.00088 | 0.0016 | NA | **0.544** |
| developmental mode | binary | 0.73 | 0.19 | 0.05 | **0.456** | **0.2534** |

### Selection for lecithotrophy in low and high salinity

Selection for lecithotrophic egg masses across three generations resulted in a proportional increase in the number of lecithotrophic egg masses in both low and high salinity (Figure 3, Table 2). As the sample size for the low salinity S_2_ generation was very small (1 family line, 4 individuals) only the S_2_ generation for the high salinity treatment is shown (three family lines, 17 individuals). The response to selection was similar for both low and high salinity selected lines, while the selection coefficient was greater for the low salinity selected lines (Table S1). Selection increased the proportion of mixed egg masses in both low and high salinities (Table 2, Figure 3). The summed realized heritability for egg capsule size was 0.39 for high salinity and 0.34 for low salinity (Table 2). Similarly, for developmental mode realized heritability was 0.35 for high salinity and 0.38 for low salinity (Table 2, Table S1). Changes in the slope of the reaction norm following selection was tested with 349 individuals from 9 families (Long Beach: n= 5, Mill Valley: n=4) selected in both low and high salinity. The slope of the reaction norm remained positive following selection (Figure 4), and a linear model with the response proportion lecithotrophy against the additive predictors salinity and generation was not statistically significant (salinity p-value = 0.21, generation p-value 0.30, r^2^ 0.12, p-value = 0.25).

**Table 2.**
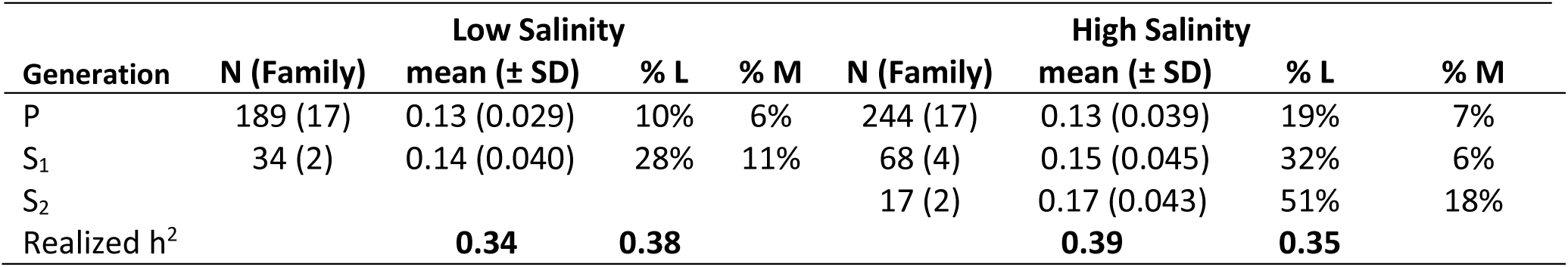
Selection for lecithotrophy in low (16 ppt) and high (32 ppt) salinity showing the mean egg capsule size and the proportion of lecithotrophic egg masses (prop. l), proportion mixed egg masses (prop. m), and their realized heritabilities. N is the number of individuals that survived to lay eggs. The number of maternal family lines is parenthetical to the number of individuals. The data for the parental generation is pooled from two separate experiments.

**Figure 3.**
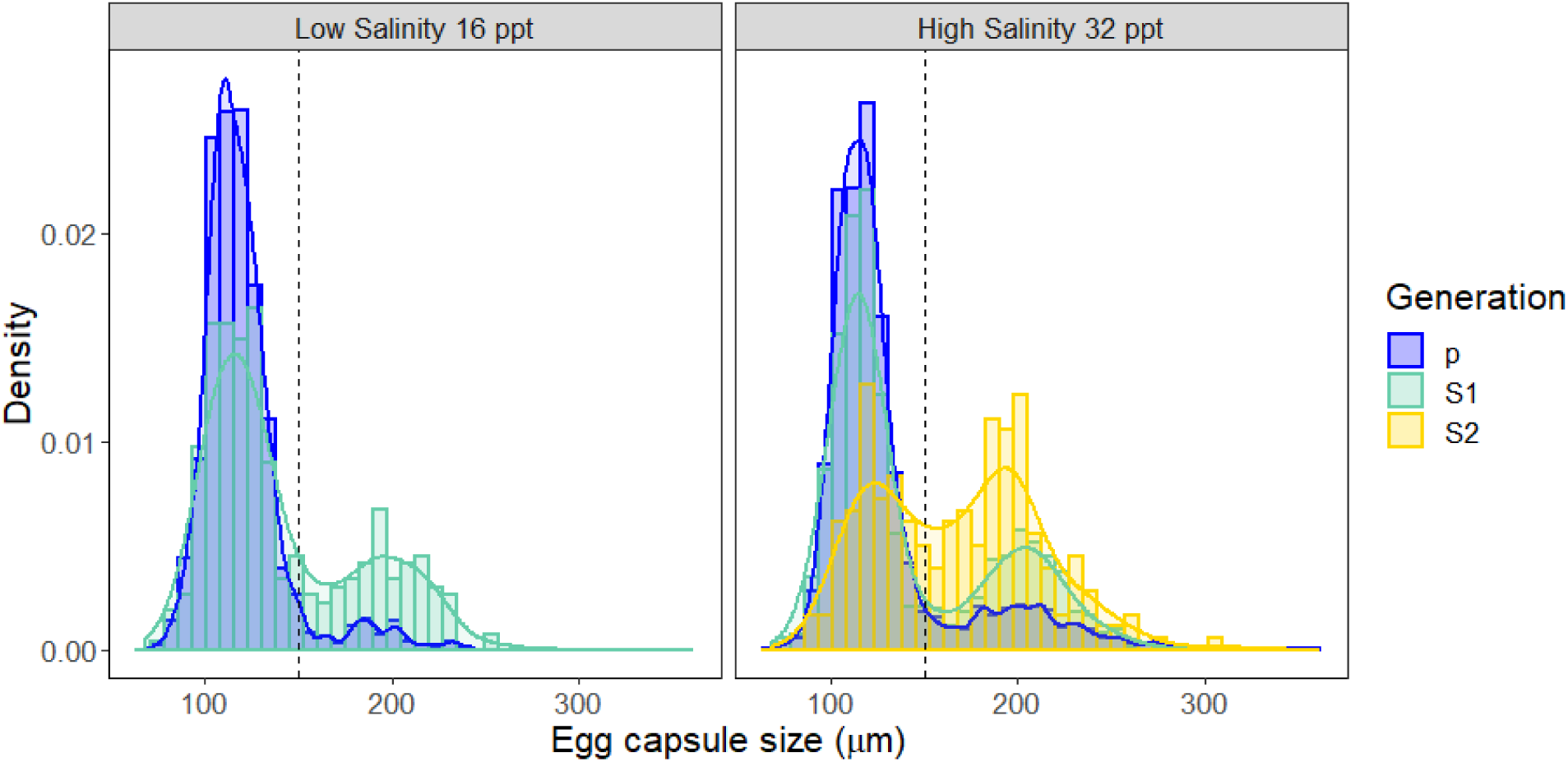
Barplot with density overlay showing the response to selection for lecithotrophy across several generations in low (16 ppt) and high (32 ppt) salinity. The vertical dashed line indicates the cut-off for lecithotrophic or planktotrophic development (egg capsule size 150 µm). Generations S_1_ and S_2_ are ‘selfed’ (see Methods) while the parental generation is the product of outcrossing in the field. The S_2_ generation in low salinity is not shown due to small sample size (4 individuals from a single family, all of which laid lecithotrophic egg masses).

**Figure 4.**
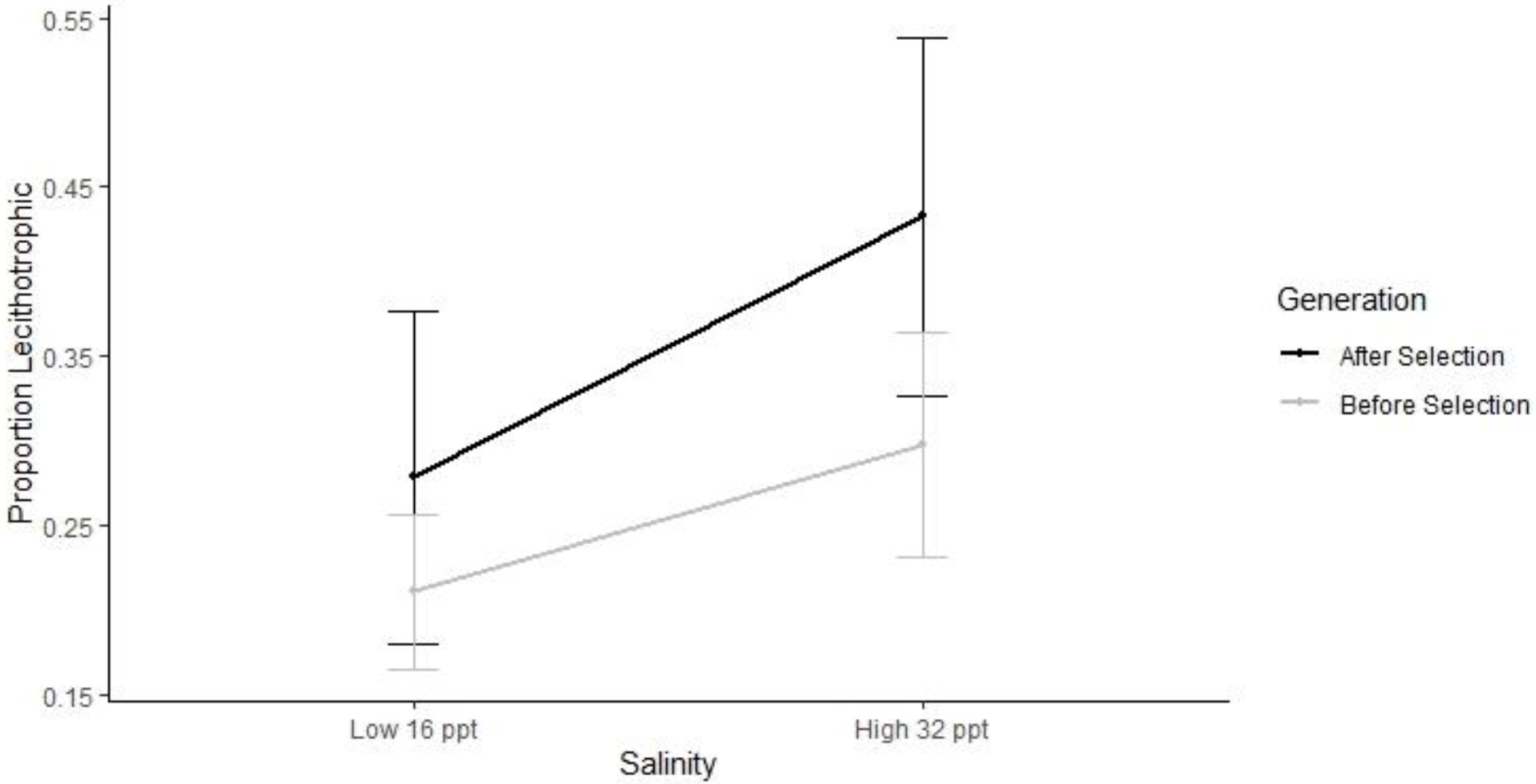
The reaction norm before and after one generation of selection for lecithotrophy. Note that this is a subset of the data (9 Families, 5 Long Beach, 4 Mill Valley) presented in Table 2 as not all families laid enough egg-masses to be split into low and high salinity following selection (See methods and results).

## Discussion

The sacoglossan sea slug *Alderia willowi* exhibits variation in egg size leading to two developmental modes, lecithotrophy and planktotrophy, with differing developmental durations and dispersal potentials. Previous studies have shown that in *A. willowi* intraspecific variation in developmental mode (poecilogony) is a seasonal polyphenism modulated by the environment experienced by juvenile slugs (Krug et al. 2012). This study confirms experimentally that variation in the production of planktotrophic vs. lecithotrophic offspring is at least partly conditional on ambient salinity, but that the response varies across families, indicating a strong genotype by environment interaction. Developmental mode in *A. willowi* responds readily to selection for increased proportions of lecithotrophy. The response to selection implies sufficient genetic variation for developmental mode and has implications on both the phylogenetic rarity of poecilogony as well as the maintenance of plasticity in *A. willowi* as slug populations appear capable of quickly responding to seasonally shifting environmental conditions.

Salinity is a common stressor for estuarine animals that varies seasonally in California (Cloern et al. 2017). Most maternal families of *A. willowi* responded to low salinity by producing more planktotrophic offspring, mirroring the pattern found in natural populations during the winter months when seasonal rain lowers mean and minimum salinity and proportions of planktotrophic egg masses increase (Patrick J. Krug, Gordon, and Romero 2012). These results suggest salinity is a cue for the type of egg mass to lay as well as a source of stress, as low salinity led to reduced survival, particularly in self-fertilized sibships (see Table 2). Seasonally fluctuating selection pressures are well documented in several taxa and represent challenging conditions to which non-migratory organisms must adapt through plasticity or, for organisms with short generation times, through changes in allele frequency, or via a combination of plasticity and allele frequency change (Bergland et al. 2014; Jones and Robinson 2018; Kingsolver and Buckley 2017).

Plasticity for developmental mode is exceptionally rare and has only been documented in *A. willowi* (Krug et al. 2007) allowing for the examination of environmental as well as genetic factors that influence the evolution of developmental mode. Studies of poecilogonous species have been critical to furthering our understanding of developmental mode evolution (Levin et al. 1991, Zakas and Wares 2012, Zakas and Rockman 2014). In particular, genetic crosses between individuals with lecithotrophic and planktotrophic development in the poecilogonous polychaete annelid *Streblospio benedicti* have revealed that developmental mode is highly modular with several large-effect loci that influence maternally determined egg size and other loci that act independently to determine larval morphology (e.g., chaeta length, Zakas et al. 2018). Yet in sacoglossan sea slugs, egg size variation appears to be due to many small effect loci, as shown by crosses between lecithotrophic and planktotrophic individuals in the poecilogonous sea slug *Elysia chlorotica* (West H.H, Harrigan J.F., Pierce S.K. 1984) and the results presented herein *for Alderia willowi* with the addition of environmental sensitivity. In both annelids and sacoglossan there is support for a step-wise model of the evolution of developmental mode, but based on the handful of species to be thoroughly examined they vary in the number of loci and their effect size. These differing patterns of inheritance between annelids and mollusks suggest that alternate genetic pathways have contributed to developmental mode between these two phyla. Additional case studies that perform genetic crosses, QTL mapping, and genomic or transcriptomic sequencing would determine whether these genetic pathways are novel or shared between genera or phyla.

## Supplemental tables and figures

**Supplementary Table S1.** Parameters for realized heritability in low and high salinity for egg capsule size and the proportion of lecithotrophy. Where q is the proportion of lecithotrophy pre-selection when applicable, µ is the mean trait value, S is the selection coefficient, R is the response, and H^2^ is the broad sense heritability. Sum H^2^ is calculated by dividing the summed responses by the summed selection coeffiecients for each trait and salinity separately.

| Proportion<br>Lecithotrophic |  | Low salinity |  |  |  |  | High salinity |  |  |  |  |
| --- | --- | --- | --- | --- | --- | --- | --- | --- | --- | --- | --- |
| Generation | $q$ | $u$ | $s$ | $R$ | $H^2$ | | $q$ | $u$ | $S$ | $R$ | $H^2$ |
| P | 0.10 | -1.28 | 1.75 | 0.70 | 0.40 |  | 0.19 | -0.88 | 1.43 | 0.41 | 0.29 |
| S1 | 0.28 | -0.55 |  |  |  |  | 0.32 | -0.47 | 1.12 | 0.49 | 0.44 |
| S2 |  |  |  |  |  |  | 0.51 | 0.03 |  |  |  |
| sum $H^2$ | | | | | | | | | | | 0.35 |
| Egg capsule size |  |  |  |  |  |  |  |  |  |  |  |
| Generation | | $u$ | $S$ | $R$ | $H^2$ | | | $u$ | $S$ | $R$ | $H^2$ |
| P |  | 0.13 | 0.06 | 0.02 | 0.34 |  |  | 0.13 | 0.05 | 0.01 | 0.24 |
| S1 |  | 0.14 |  |  |  |  |  | 0.15 | 0.03 | 0.02 | 0.63 |
| S2 |  |  |  |  |  |  |  | 0.17 |  |  |  |
| sum $H^2$ | | | | | | | | | | | 0.39 |

**Figure S1.**
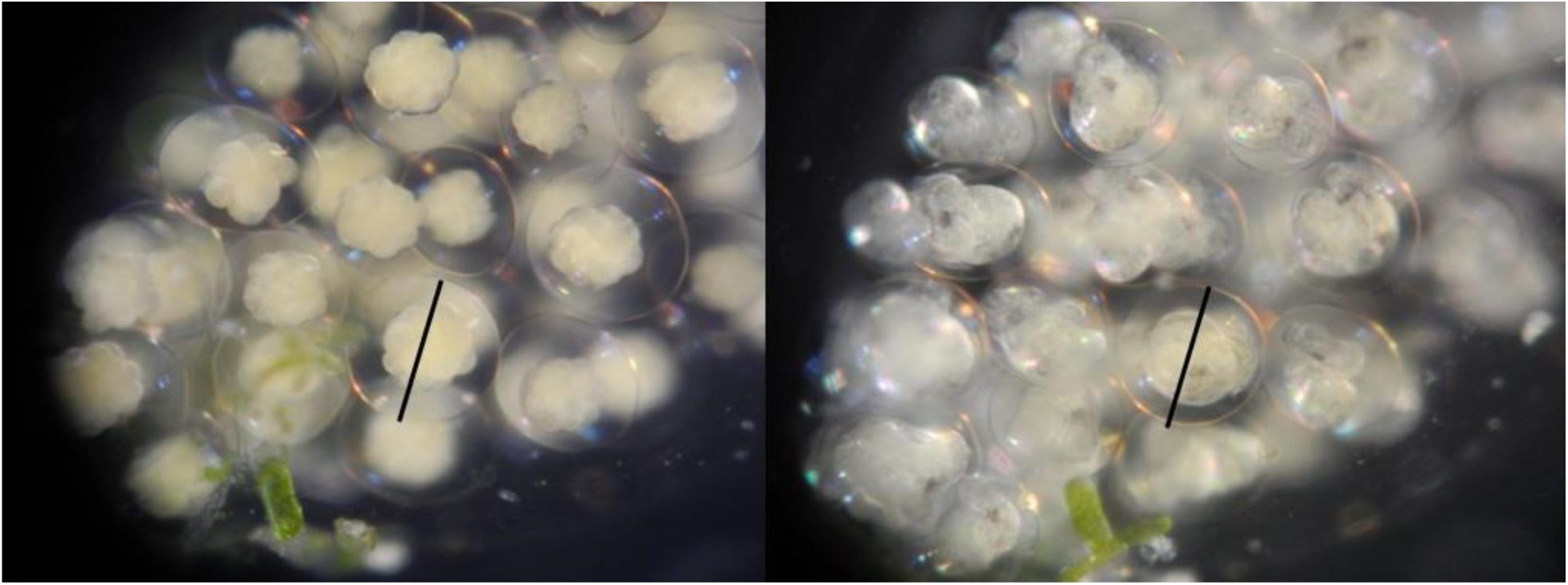
Egg capsule size remains constant across larval development as seen in these figures showing the same embryo at the 32-64 cell stage and 4 days later at the veliger stage. At each time the egg capsule was measured to be 206 µm.

**Figure S2.**
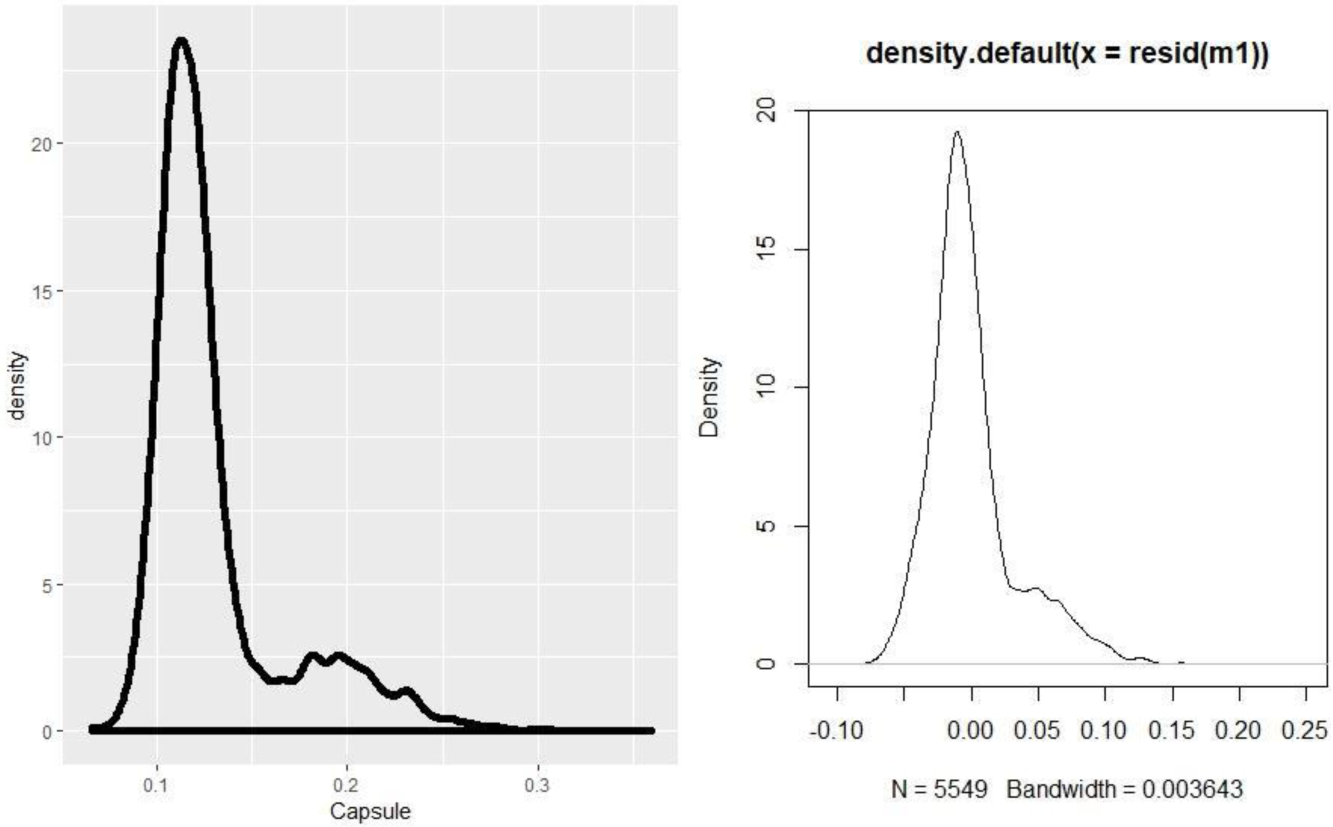
Non-normality of raw data (egg capsule size) and residual in a linear model where egg capsule size is the response and Family and Salinity are the predictors. Normality can be achieved in the residuals if developmental mode (lecithotrophy versus planktotrophy) based on an egg capsule size cutoff of 150 µm is used.

## Notes

https://github.com/SerenaCaplins/GXE_A.willowi

